# Uncovering the architecture of selection in two *Bos taurus* cattle breeds

**DOI:** 10.1101/2021.11.11.468293

**Authors:** Troy N. Rowan, Robert D. Schnabel, Jared E. Decker

## Abstract

Selection alters the genome via hard sweeps, soft sweeps, and polygenic selection. However, mapping polygenic selection is difficult because it does not leave clear signatures on the genome like a selective sweep. In populations with temporally-stratified genotypes, the Generation Proxy Selection Mapping (GPSM) method identifies variants associated with generation number (or appropriate proxy) and thus variants undergoing directional allele frequency changes. Here, we use GPSM on two large datasets of beef cattle to detect associations between an animal’s generation and 11 million imputed SNPs. Using these datasets with high power and dense mapping resolution, GPSM detected a total of 294 unique loci actively under selection in two cattle breeds. We observed that GPSM has a high power to detect selection in the very recent past (< 10 years), even when allele frequency changes are small. Variants identified by GPSM reside in genomic regions associated with known breed characteristics, such as fertility and maternal ability in Red Angus and carcass merit and coat color in Simmental. Over 60% of the selected loci reside in or near (<50 kb) annotated genes. Additionally, 36% of selected loci overlap known epigenetic marks or putative functional genomic regions. Using RAiSD and nSL, we identify hundreds of putative selective sweeps; however, these sweeps have little overlap with polygenic selected loci. This makes GPSM a complementary approach to sweep detection methods when temporal genotype data are available. The selected loci that we identify across methods demonstrate the complex architecture of selection in domesticated cattle.

## Introduction

Since their initial domestication in the Fertile Crescent ~10,500 years ago (Bradley et al. 1996) cattle have been exposed to intense selection for increased tameness, production, and fecundity leading to substantial changes in these phenotypes. While selection occurs on phenotypes and very recently on estimated breeding values, the phenotypic changes of a population are due to changes in the genotype frequencies of the trait’s underlying causal variants. Occasionally, selection on beneficial mutations of substantial phenotypic effect can lead to rapid allele frequency changes at a locus. The resulting “selective sweep” not only increases the frequency of the beneficial variant but also reduces genetic variation in the genomic region surrounding the selected locus (Smith and Haigh 1974). In cattle populations, multiple sweeps have been mapped to genomic regions, many of which are related to simple Mendelian traits such as polledness (the absence of horns) (Ramey et al. 2013) and coat color (Boitard et al. 2016) or large-effect quantitative trait loci (QTL) influencing production traits (Gutiérrez-Gil et al. 2015).

While sweeps at Mendelian loci have played an essential role in the domestication and subsequent improvement of cattle populations, it is becoming increasingly apparent that the majority of both historical and ongoing selection is on highly complex traits (Kemper et al. 2014; Zhang et al. 2020). Complex traits are controlled by many mutations of relatively small effect spread throughout the genome (Mackay et al. 2009). Under complex trait architectures, selection can generate substantial changes to a phenotype without necessarily generating large allele frequency shifts (Barghi et al. 2020). These modest directional allele frequency changes at selected loci make mapping selection on polygenic traits over short timescales difficult. However, by leveraging large commercially-generated datasets that include influential founder individuals, cattle populations offer intriguing opportunities for mapping the loci exposed to polygenic selection over time (Decker 2015). The Generation Proxy Selection Mapping (GPSM) method uses a proxy for the number of meioses that separate an individual from the beginning of the pedigree as the dependent variable in a genome-wide linear mixed model to detect significant associations between generation and allele frequency caused by selection (Decker et al. 2012; Walsh and Lynch 2018). When pedigrees have missing data or complex, overlapping generations, a proxy for generation number, such as birth date, can improve the analysis. Both simulations and empirical data show that GPSM effectively detects subtle ongoing shifts in allele frequency across the genome of multiple populations (Rowan et al. 2021).

Genome-wide association studies have motivated extensive work attempting to dissect the genetic architecture of complex traits in many species (Timpson et al. 2018). We expect that selection on these complex traits occurs on similar polygenic architectures (Barghi et al. 2020). Here, we expand on previous work exploring the selection landscape in cattle with one of the largest non-human selection mapping datasets analyzed to date. Using sequence-imputed genotypes for over 124,000 individuals in two beef cattle populations, we identify subtle ongoing shifts in allele frequency due to selection at over 11 million sequence variants. With these high-resolution data, we can discern the underlying functional genomics on which selection acts in cattle populations. Further, knowledge of ongoing selection in the bovine genome can serve as important annotations that add context to genomic regions identified in other studies. Additionally, we use subsets of these data to explore recent and ongoing selective sweeps using both haplotype and site frequency spectrum (SFS)-based approaches. This combination of selection mapping approaches provides a detailed report of the genetic architectures on which strong directional selection acts in cattle populations and a blueprint for leveraging temporally distributed genotypes to understand the selection architectures of other species.

## Results

### Quantitative Genetic Signals of Polygenic Selection

In each population, we used genomic restricted maximum likelihood (GREML) (Yang et al. 2010) to estimate the proportion of variance explained (PVE) by 811,967 imputed SNPs in various subsets of our Simmental (SIM) and Red Angus (RAN) datasets (**Supplementary Tables 1 & 2**). Using an individual’s date of birth as a generation proxy, we estimated the PVE to be 0.523 (SE = 0.007) and 0.619 (SE = 0.005) in RAN (n = 46,454) and SIM (n = 78,787), respectively. Results from simulations in our previous work suggest that PVE is mostly a function of the demographics of a population and the strength of selection occurring (Rowan et al. 2021). Due to the non-normality of sampled birth dates in both datasets (**Supplementary Figure 1**), we observe a large divergence from expectation in individual breeding values and residuals in GREML analyses, particularly for the earliest born individuals (**Supplementary Figure 2**). Although linear mixed models are robust to model misspecification (Visscher et al. 1996; Zhou et al. 2013), we explored transformations to increase our power. We performed a Box-Cox transformation to birth date to help normalize residuals and potentially boost our power to detect selected variants (**Supplementary Tables 1 & 2**). When using Box-Cox-transformed birth date as the dependent variable in Red Angus, PVE increased to 0.657 (S.E. = 0.006). A Box-Cox transformation to birth date in Simmental decreased PVE to 0.605 (SE = 0.005). This transformation noticeably normalized GREML-estimated residuals and breeding values in both datasets (**Supplementary Figure 3**).

We explored the effects of various statistical transformations on our ability to detect GPSM signatures in Red Angus using the 811K SNP dataset. The number of significant SNPs identified using an identical dataset, but with different statistical transformations performed on the generation proxy provides a measure of statistical power. In Red Angus, we observed an increase in the number of variants detected at multiple significance thresholds (nominal: p < 10^−5^, Bonferroni: p < 7.55×10^−7^, and FDR-corrected q-values < 0.1 and < 0.05). At all significance thresholds, Box-Cox transformed birth date detected between 71% to 89% more significant SNPs and at least 12 additional significant loci (**Supplementary Table 3**). Interestingly when analyzing data only for animals born since 2012, GPSM identified almost all of the same loci as the Box-Cox transformed birth date GPSM on the full Red Angus dataset. The same transformation applied to the Simmental dataset reduced the number of identified SNPs and loci compared with raw birth date (**Supplementary Table 4**).

While the Red Angus dataset was composed of almost entirely purebred individuals, the Simmental dataset contained large numbers of crossbred animals. The numbers of animals with non-Simmental ancestry that have been genotyped by the breed association have significantly increased in recent generations (**Supplementary Figure 4**). Consequently, we divided the Simmental dataset into subsets based on pedigree-reported ancestry and birth date. This allowed us to examine the selection that is occurring within different subsets of the population. The PVE for birth date remained moderately high, ranging from 0.619 (SE = 0.005) for all animals with > 5% SIM ancestry to 0.436 (SE = 0.021) for animals born before 2008. A complete accounting of variance components estimated from these subsets is in **Supplementary Table 2**.

### Generation Proxy Selection Mapping in Red Angus

Based on results from the 811K GPSM analyses described above, in Red Angus using birth date as the dependent variable, we performed three separate sequence-level analyses (11,759,568 imputed SNPs) for all individuals, for animals born before 2012, and animals born after 2012, referred to hereafter as full Red Angus, old Red Angus, and young Red Angus, respectively.

GPSM identified 2,914 SNPs, 9,065 SNPs, and 0 SNPs (Bonferroni-corrected threshold p < 4.29 × 10^−9^) significantly associated with birth date in the full, young, and old Red Angus datasets, respectively (**Figure 1**). A less stringent p-value threshold of 5 × 10^−8^ identified 3,617, 10,939, and 0 SNPs associated with birth date in the full, young, and old Red Angus datasets, respectively. The old dataset, which contained only 1,984 individuals, was likely underpowered to detect signals of selection using GPSM. The full dataset identified 3,240 SNPs that were also significant in the young Red Angus data, a near-complete overlap. A stepwise conditional & joint analysis (COJO) (Yang et al. 2012) of these results further refined lead SNPs in significant loci (GPSM and COJO p < 5×10^−8^). COJO identified 72 genome-wide significant (p < 5 × 10^−8^) independent associations with birth date in the full dataset and 96 in the young dataset. Despite these datasets being composed mainly of the same individuals and sharing a large proportion of common GPSM SNPs, only 16 SNPs identified by COJO were shared between both datasets.

**Figure 1.**
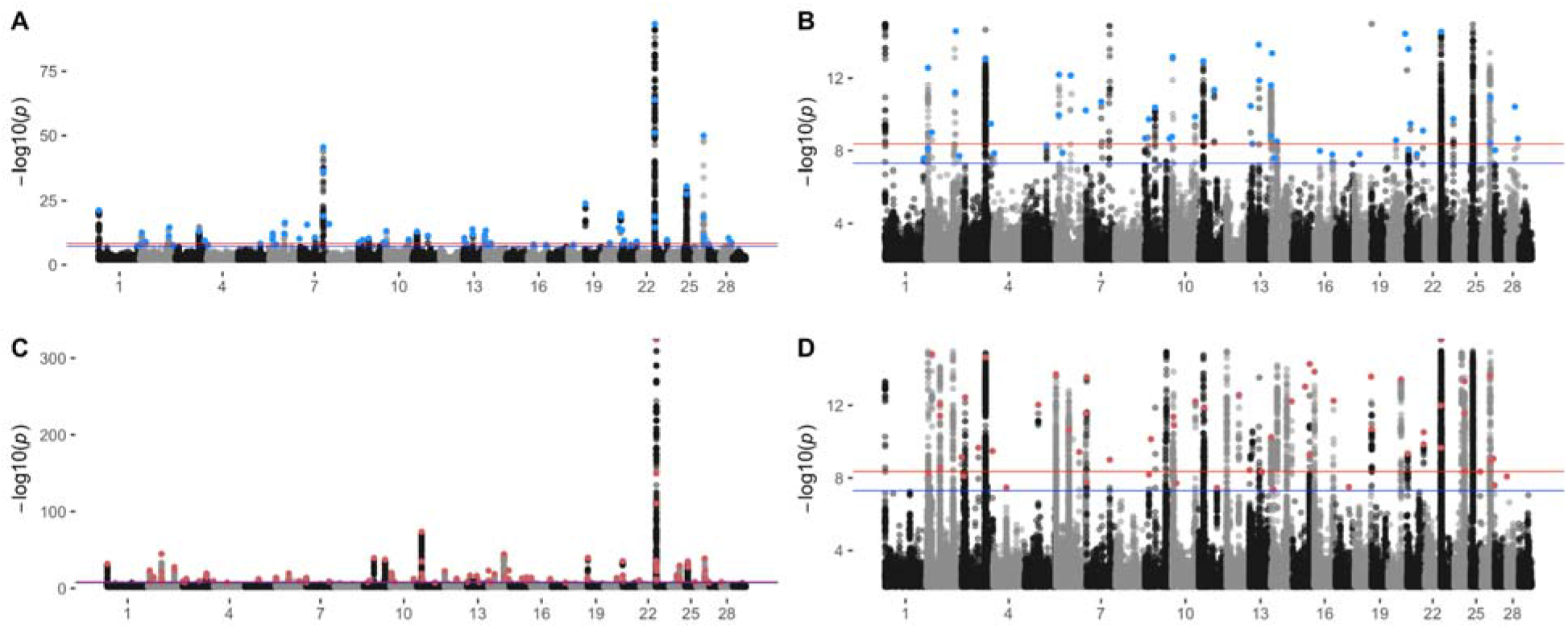
GPSM detects polygenic selection in the Red Angus dataset. Red Angus GPSM Manhattan plots for the (A) full dataset, (B) truncated at p = 10^−5,^ and (C) young dataset also truncated (D) at p = 10^−5^. Genome-wide significance is indicated by the red line (Bonferroni-corrected p-value = 4.29 × 10^−9^) and blue line (p-value = 5 × 10^−8^). Blue points are significant COJO SNPs (COJO p < 5 × 10^−5^) in the full Red Angus dataset. Red points are significant COJO SNPs in young Red Angus datasets.

Using COJO SNPs from the sequence-level GPSM analysis, we annotated nearby genes, known quantitative trait loci (QTL), and other genomic features to help understand the biological pathways and phenotypes that selection targets in this population. In all datasets, the majority (51-72%) of COJO SNPs resided within or adjacent to (within 50 kb) annotated genes (**Table 1**). Depending on the dataset, 40% (full) or 54% (young) of these COJO SNPs with a nearby gene resided in regions immediately upstream or downstream of transcription start sites, insinuating that selection is primarily acting on *cis*-regulatory regions of the genome. A complete accounting of nearby genes for COJO SNPs in the full and young Red Angus datasets is provided in **Supplementary Tables 5 & 6**, respectively.

**Table 1.** Number of significant (COJO and genome-wide p < 5 × 10^−8^) SNPs within, proximal, or outside genic regions.

| Breed | Dataset | Total SNPs | COJO | Within Gene | Gene Proximal (< 50kb to nearest) | Intergenic |
| --- | --- | --- | --- | --- | --- | --- |
| Red Angus | Full | 72 |  | 17 (24%) | 20 (28%) | 35 (48%) |
| Red Angus | Young | 96 |  | 36 (38%) | 24 (25%) | 36 (38%) |
| Simmental | Full | 108 |  | 32 (30%) | 36 (33%) | 40 (37%) |
| Simmental | Purebred | 18 |  | 6 (33%) | 7 (39%) | 3 (17%) |

In many cases, GPSM in young Red Angus animals detects novel signatures of recent ongoing selection that are not significant in the full dataset. Chromosome 2 offers an interesting case study of these differences. In addition to the two significant peaks identified by both the full and young Red Angus datasets, at least eight additional major peaks are identified in the young data, accounting for 48 unique COJO associations (**Figure 2 A & B**). The strongest unique associations identified in the young Red Angus dataset reside within the gene *ARHGAP15*. This association contains four unique COJO SNPs within the gene. *ARHGAP15* is a major gene involved in trypanotolerance in African cattle (Noyes et al. 2011; Álvarez et al. 2016) and likely has broader effects on immune function in worldwide cattle populations. Further, *ARHGAP15* is almost exclusively expressed in immune tissues (Fang et al. 2020).

**Figure 2.**
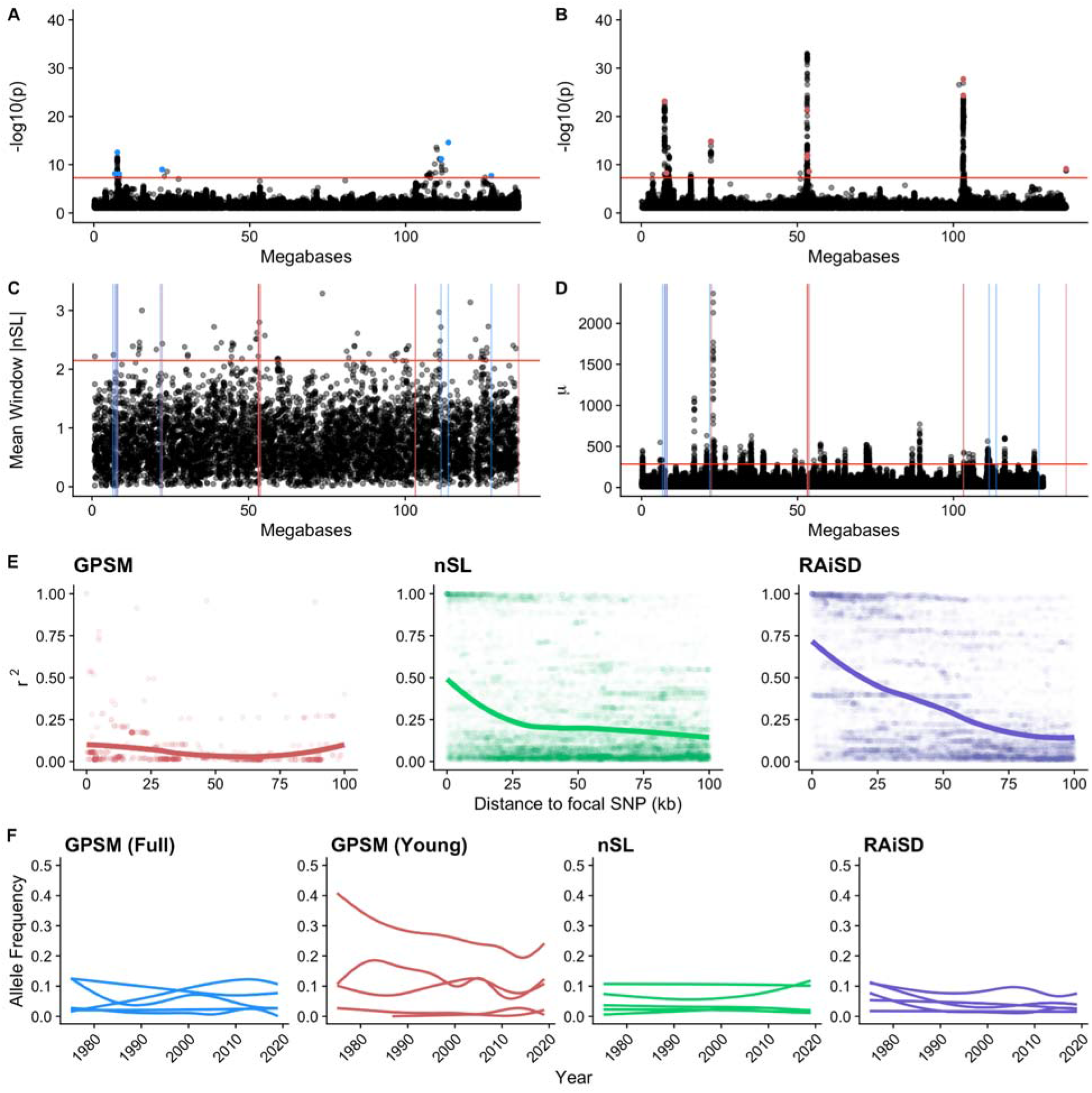
Methods identify largely different regions of selection on chromosome 2 in Red Angus. Chromosome 2 Manhattan plots for (A) full and (B) young Red Angus datasets. The red line is a Bonferroni-corrected significance threshold (4.29 × 10^−9^), and the blue line is a less stringent genome-wide significance threshold of 5 × 10^−8^. (C) Genomic distribution of |nSL|scores for windows defined b GenWin and (D) RAiSD μ statistics. Horizontal red lines indicate 0.05% outlier genome-wide thresholds. Vertical red and blue lines represent the positions of full Red Angus and young Red Angus GPSM COJO SNPs, respectively. (E) Pairwise linkage disequilibrium (r^2^) values for SNPs within 100 kb of COJO SNPs for GPSM, SNPs closest to the center of significant nSL windows, or lead SNPs in significant RAiSD peaks. (F) Allele frequency trajectories over time for the five most significant SNPs in each analysis.

Previously identified and annotated QTL for cattle allowed us to interpret the biological and production impact of selection decisions reflected in the GPSM results. Using QTL annotations, we can extrapolate the combined selective pressures on a population without ascribing a particular genomic feature to a selected locus (e.g., a mutation within a transcription factor binding site). A QTL-enrichment analysis identified 5 QTL classes significantly enriched (FDR-adjusted p < 0.05) within 50 kb of GPSM COJO associations (**Supplementary Tables 7 & 8**). The most significant QTL enrichment in Red Angus was for metabolic body weight (FDR-corrected p = 8.19 × 10^−96^), driven primarily by GPSM signals on chromosomes 14 (23.3 Mb) and 6 (38.5 Mb). Other enriched QTL pointed towards ongoing selection for increased fertility and calving ease, known selection goals in the breed (https://redangus.org/about-red-angus/history/). Calving ease, sexual precocity, and carcass weight QTL were each enriched among QTL tagged by GPSM COJO SNPs across the genome (FDR-corrected p < 4.15 × 10^−3^). We did not observe any enriched gene ontologies or pathways in the full Red Angus GPSM analysis. Twelve terms were enriched in the young Red Angus GPSM results (**Supplementary Table 9**), driven mainly by three genes (ENSBTAG00000049666/*CYP3A5*, ENSBTAG00000052665/*CYP3A4*, ENSBTAG00000053645/*CYP3A5*).

### Generation Proxy Selection Mapping in Simmental

We analyzed our dataset’s purebred Simmentals (greater than 87.5% Simmental pedigree ancestry) to identify ongoing selection restricted to the breed. We performed an additional sequence-level GPSM on the full Simmental dataset (all animals with ≥ 5% reported Simmental ancestry), representing selection within the entire herdbook, including the full spectrum of admixed animals.

GPSM analysis of purebred Simmental animals (n = 13,379) identified 513 genome-wide significant SNPs (**Figure 3 A & B**) (Bonferroni-corrected p-value threshold p < 4.29 × 10^−9^). A more relaxed significance threshold (p < 5 × 10^−8^) identified 642 birth year-associated SNPs. A strong signal on chromosome 5, centered at a locus containing *PMEL* and *ERBB3*, genes is known to control coat color in Fleckvieh populations (i.e., European Simmental) (Mészáros et al. 2015). This locus, coupled with a GPSM signature immediately upstream of *KIT* (Durkin et al. 2012), suggests that the strongest selection pressures in the purebred American Simmental population have changed coat color and external appearance. These changes have made them appear less like European Simmental and more like American Angus. The next most significant locus resides in a cluster of olfactory receptors on chromosome 28. Another strong signal exists on chromosome 15 near beta-carotene oxygenase 2 (*BCO2*) and interleukin-18 (*IL-18*), a locus likely involved in immune functions (He et al. 2010). Of these SNPs, COJO identified 18 independent associations, 2 of which were at the center of chromosome 5 in the *PMEL/ERBB3* locus.

**Figure 3.**
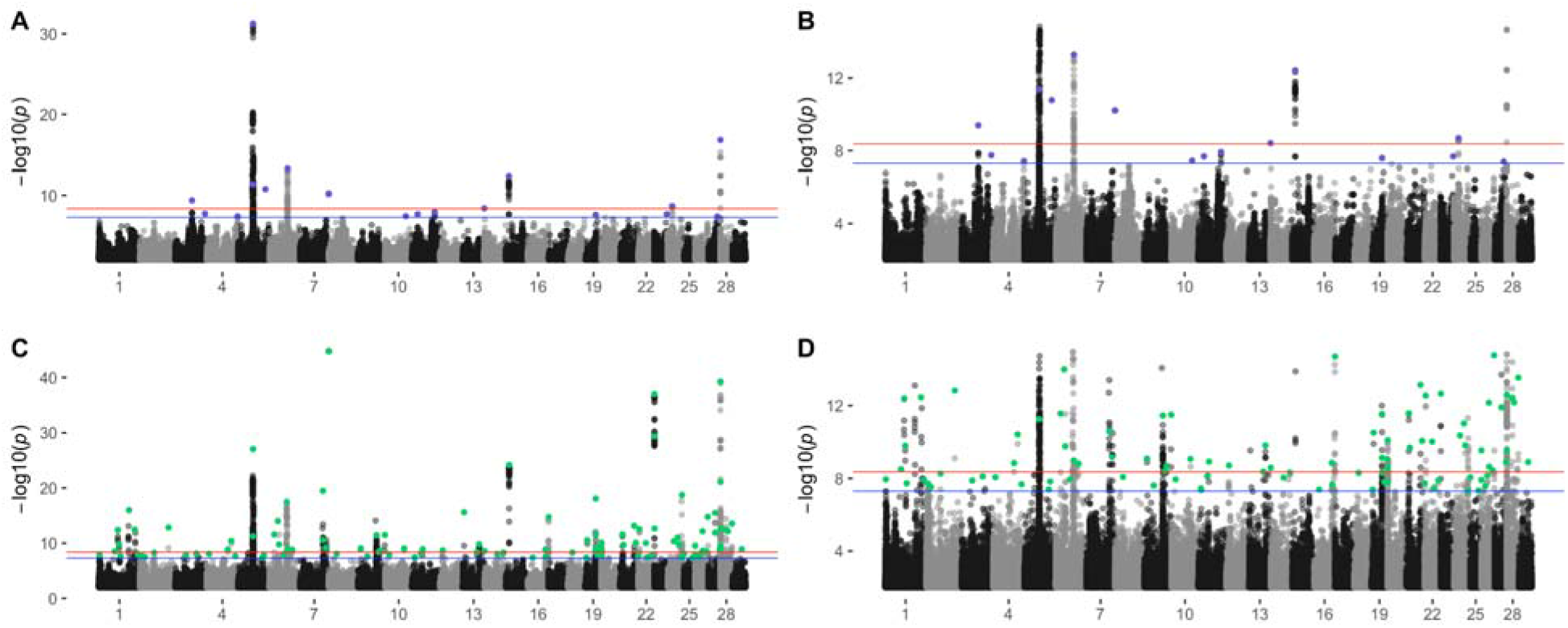
GPSM detects signatures of ongoing polygenic selection in the Simmental cattle. GPSM Manhattan plots for the (A) purebred Simmental dataset, (B) truncated at p = 10^−5^ and (C) full Simmental dataset also truncated (D) at p = 10^−5^. Genome-wide significance is indicated by the red line (Bonferroni-corrected p-value = 4.29 × 10^−9^) and blue line (p-value = 5 × 10^−8^). Purple points indicate significant COJO SNPs (COJO p < 5 × 10^−5^) in the purebred Simmental dataset. Green points are significant COJO SNPs in the full Simmental dataset.

A GPSM analysis of all registered Simmental animals with at least 5% Simmental pedigree ancestry (n = 78,787) identified 1,008 genome-wide significant SNPs (Bonferroni-corrected p < 2.29 × 10^−9^), and 1,356 significant SNPs at a slightly less strict threshold (**Figure 3 A & B**). A COJO analysis found 108 independently-associated SNPs (GPSM and COJO p < 5 × 10^−8^), two of which were also identified in the purebred analysis (14:28004:G:T, 8:61247:T:C). These include the same significant coat color-associated regions on chromosomes 5 and 6 and olfactory receptor clusters on chromosome 28 mentioned above.

By subsetting these data, we identified recent Simmental-specific selection in the purebred dataset. We contrasted it with signatures identified in the full dataset, where introgression from other breeds is also responsible for changing allele frequencies. In the full dataset, the most significant COJO SNP ( 9:85445646:T:C, COJO p = 2.51 10^−276^) is located immediately upstream of *SASH1*, a gene implicated in various fertility phenotypes in beef cattle (Fortes et al. 2010; Xiang et al. 2017; Sweett et al. 2020). We also identified strong selection on variants (4:94308044:A:G, COJO p = 2.17 × 10^−170^) in the imprinted gene *COPG2* (Khatib et al. 2007). As in Red Angus, the majority of GPSM COJO SNPs were either in or proximal to (< 50 kb from) genes (**Table 1**). 29% of the full and 33% of purebred COJO SNPs resided within genes. An additional 33% and 39% of COJO SNPs resided close to genes in the full and purebred datasets, respectively. A complete accounting of Simmental GPSM COJO detected SNPs and their nearby genes are in **Supplementary Tables 10 & 11.**

In the purebred Simmental dataset, two of the most significant QTL enrichments were involved with the appearance-based traits eye pigmentation (FDR-corrected p = 4.24 × 10^−5^) and coat color (FDR-corrected p = 4.29 × 10^−1^), primarily driven by the selected loci on chromosomes 5 and 6 (**Supplementary Table 12**). While these same significant loci were also identified in the full Simmental GPSM analysis, the enriched QTL classes under selection in the full dataset also included carcass (longissimus muscle area and carcass weight), production (metabolic body weight, average daily gain, dry matter intake), and reproduction (Inhibin level, luteal activity) (**Supplementary Table 13**). Both Simmental datasets identified enrichments in keratinization gene sets (**Supplementary Tables 14 & 15**).

### Ongoing selection at annotated loci

The Functional Annotation of Animal Genomes (FAANG) project has generated genome-wide data for multiple epigenetic marks and open chromatin (CTCF, H3K4me1, H3K4me3, H3K27me3, H3K27ac, and ATAC-seq) in multiple tissues (adipose, cerebellum, cortex, hypothalamus, liver, lung, muscle and spleen) in a pair of biological replicates (Giuffra et al. 2019). These data, coupled with dense GPSM results, allowed us to understand how selected loci overlap with annotated epigenetic and open chromatin marks of the genome. Previous selection mapping studies could generally resolve significant loci to nearby genes. GPSM combined with COJO allowed us to refine the selected variant to very small regions (SNPs approximately every 250 bp) driving the signal. Twenty-six COJO SNPs (36%) from the full Red Angus dataset resided in at least one of these epigenetic classes in at least one tissue. In the young Red Angus dataset, 35 of the COJO SNPs resided in one of these regions (36%). GPSM signal is not significantly enriched in any single epigenetic mark region (Binomial test p-values all > 0.39). That said, some GPSM SNPs may represent selection on regulatory variation. These FAANG annotations allow us to more thoroughly categorize some of the intergenic variation that we observe under selection. For example, a significant COJO SNP on chromosome 26 at 49,432,811 bp is immediately downstream of the gene Glutaredoxin-3 (*GLRX3*). The region containing this SNP harbors all six epigenetic marks, appearing in all eight tissue types across both biological replicates. Regulation of *GLRX3* has been postulated to play roles in feed efficiency (Seabury et al. 2017) and calving ease (Purfield et al. 2020). While the marks near *GLRX3* are numerous, most other regions are more specialized in their mark type and tissue context. For example, GPSM COJO identified a significant intronic SNP within the *B3GALT1* gene that resides within an H3K27ac active enhancer mark in only liver tissue. The *B3GALT1* gene is associated with feed efficiency growth traits in beef cattle (Zhang et al. 2020) and gene expression differences occur in cattle with growth retardation (Kong et al. 2020). Taken together, these results suggest that ongoing selection for growth and efficiency traits are driving allele frequency changes to regulatory variation proximal to the *B3GALT1* gene.

To determine if selected loci were more likely to play functional roles in the genome, we annotated COJO SNPs with their Functional and Evolutionary Trait Heritability (FAETH) scores as described in Xiang et al. (Xiang et al. 2019). SNPs in the top 1/3 of the FAETH score distribution (per-variant score > 1.607 × 10^−8^) were considered likely to be functional, as in Xiang et al. In Red Angus, 27% and 41% of COJO SNPs had FAETH scores in the top 1/3 of the FAETH score distribution for the full and young Red Angus datasets, respectively. A similar pattern existed in both Simmental datasets, where 31% and 28% of FAETH-annotated COJO SNPs were likely functional in the purebred and full datasets, respectively. However, across datasets, only 44% of GPSM COJO SNPs had FAETH scores. This is because FAETH scores were calculated in a different population and on a different genome assembly. As a result, we expect that an even more significant proportion of SNPs identified by GPSM are functional.

### GPSM identifies breed-specific balancing selection in KHDRBS2 regulatory regions

The most significant GPSM locus in both Red Angus datasets resided between 1 and 2 Mb on Chromosome 23 (**Figure 4A–B**). The locus contains multiple sub-associations with birth dates. In the young dataset, SNPs throughout the proximal 2 Mb of chromosome 23 are genome-wide significant. In this locus, 117 SNPs had p-values < 10^−310^, reported by GCTA as zero. This included the most significant SNP in the full dataset (23:1,768,070, p = 3.86 × 10^−94^). This SNP was also the most significant COJO SNP in the young Red Angus dataset, suggesting that it is the variant responsible for the signal at this locus in both Red Angus datasets. Interestingly, it was not the most significant COJO SNP in the full Simmental dataset. Rather, a SNP ~550 kb away (23:1,215,338, COJO p = 4.98 × 10^−238^) had the strongest association (**Figure 4C**).

**Figure 4.**
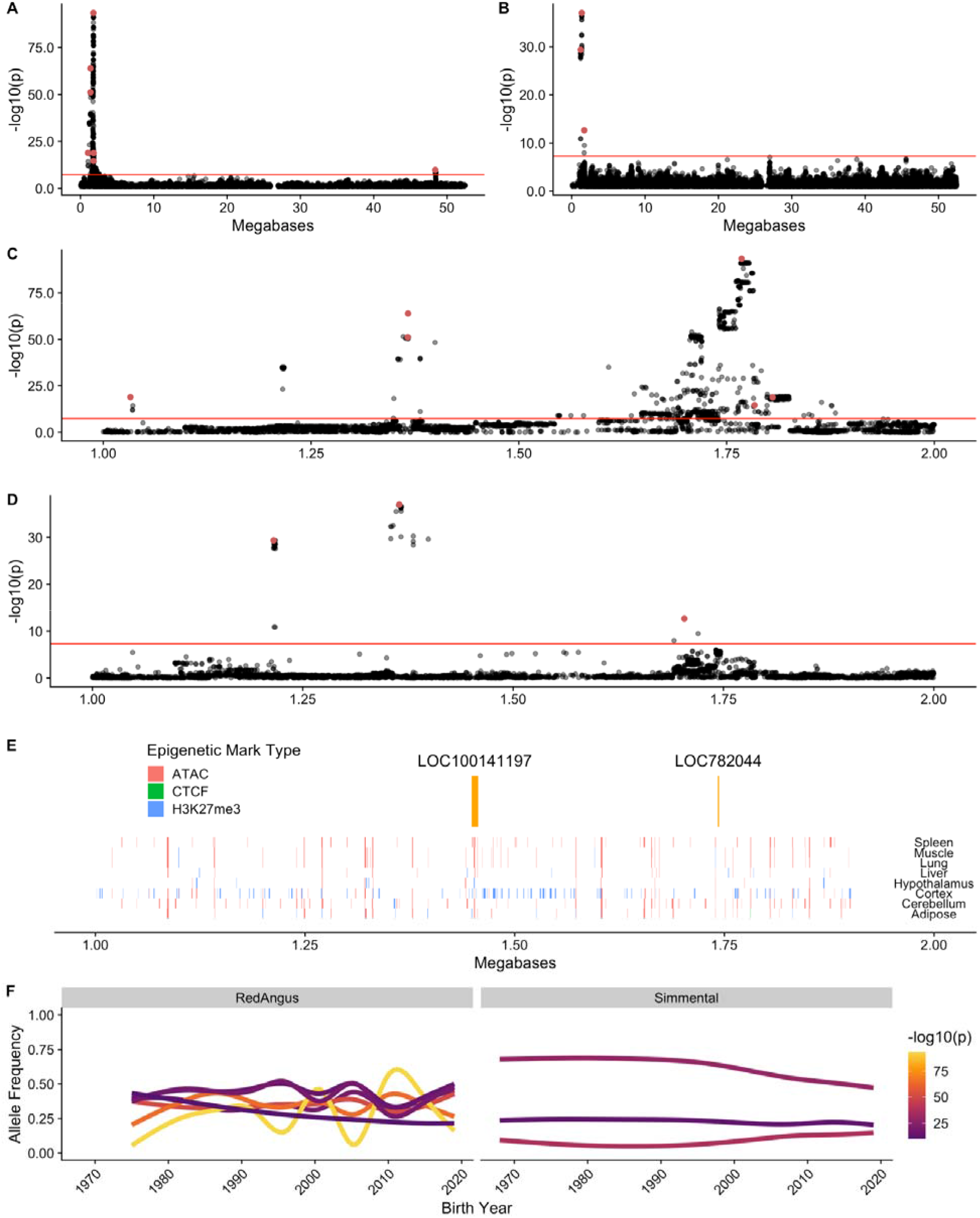
GPSM identifies a strong, complex signature of selection on chromosome 23. Manhattan plots for chromosome 23 in full (A) Red Angus and (B) Simmental datasets. Focused Manhattan plots at significant locus from 1-2 Mb on Chromosome 23 in full (C) Red Angus and (D) Simmental datasets. Red SNPs are significant GPSM COJO associations (COJO p < 5 × 10^−8^). In A-D, the red line indicates Bonferroni-corrected significance threshold. (E) Chromosome 23 (1-2 Mb) annotated with genes (orange) and epigenetic marks in eight tissues from two bovine samples in the FAANG project, colored by mark type. (F) Allele frequency trajectories represented by smoothened regression lines of the birth year versus allele frequency for significant COJO SNPs in this region in the young Red Angus and full Simmental datasets. Lines are colored by the SNP’s −log_10_(p) value from GPSM.

Forty-one SNPs within this locus were also significant in the full Simmental dataset (p < 5 × 10^−8^), including three independent COJO associations. The lead Simmental SNP at this locus resides in a secondary association identified in the young Red Angus dataset. As in Red Angus, the most significant COJO SNP (23:1,215,338:G:T, COJO p = 4.98 × 10^−238^, raw p = 4.28 × 10^−30^) was not the most significant raw GPSM p-value (23:1,364,693:A:T, raw p = 9.66 × 10^−38^). Additionally, the strongest Simmental association for this locus (between 1.2 and 1.3 Mb) differed from the strongest association identified in Red Angus (~1.7 Mb) (**Figure 4D**). This region does, however, overlap with one of the significant sub-associations found in Red Angus.

The closest protein-coding gene to this signature is *KHDRBS2*, a gene involved in reproduction in goats (Islam et al. 2020) and calving ease in cattle (Cole et al. 2011). *KHDRBS2* has also been identified as possessing a selection signature that differs between *Bos taurus* and *Bos indicus* cattle (Pérez O’Brien et al. 2014; Paim et al. 2020). While our most significant COJO SNPs do not reside directly within known epigenetic mark regions (**Figure 4E**), dozens of marks exist within this 1 Mb segment. They contain multiple genome-wide significant SNPs, suggesting strong ongoing selection for the regulation of *KHDRBS2* expression. Further, the annotated and expressed pseudogene *LOC782044* is 24,991 base pairs from Red Angus’s most significant GPSM association at 1,768,070 base pairs and 38,847 base pairs from a significant COJO SNP in Simmental. While *LOC782044* does not have any known functions in cattle, we hypothesize that it may be an enhancer RNA transcribed from an enhancer altering the expression of *KHDRBS2* or other nearby genes (Sartorelli and Lauberth 2020).

The observed allele frequencies at significant COJO SNPs in this locus show two main patterns. First, we observe small directional changes in frequency, consistent with the polygenic selection that we observe at other GPSM loci (**Figure 4 F**). Second, for some Red Angus COJO SNPs, we see allele frequencies oscillating around an intermediate value, a pattern consistent with balancing selection (Orozco-terWengel et al. 2012). Further, this region shows high Tajima’s D scores, where 25% of windows have a Tajima’s D score > 2 (top 10%), indicative of higher-than-expected levels of genetic variation. The highest Tajima’s D score genome-wide was located in a window ~70 kb from the most significant GPSM SNPs.

### Sweep mapping identifies known and novel selected loci

GPSM and traditional selective sweep methods identify allele frequency changes and their signatures on surrounding neutral sites. We used two selection mapping methods, a haplotype-based statistic, number of segregating loci (nSL)(Vatsiou et al. 2016), and a composite statistic (μ) implemented by the software RAiSD (Alachiotis and Pavlidis 2018). For nSL, we defined windows using the GenWin R package (Beissinger et al. 2015) which uses a spline function to observe changes in test statistics. We called the top 0.5% of windows significant, provided they contained at least three SNPs. Our nSL analysis in Red Angus identified 365 outlier windows on all but three chromosomes (17, 27, and 28) (**Supplementary Table 16, Supplementary Figure 5**). The correlation between nSL scores and GPSM effect sizes for 811K SNPs was −0.007. None of the significant GPSM COJO SNPs resided in outlier nSL windows. We also used RAiSD to look for site frequency spectrum differences indicative of selection. RAiSD calculates μ in 50 SNP sliding windows (mean length = 39,139 bp), and we considered windows with the top 0.05% of μ values as signatures of selection (**Supplementary Table 17**). The RAiSD analysis of sequenced Red Angus animals in the Thousand Bulls Project (n = 14) (Hayes and Daetwyler 2019) identified 3,740 significant windows, many of which were overlapping, that encompassed at least 339 loci exhibiting sweep-like signatures.

We found that 50% of selection nSL windows and 53% of selection RAiSD windows contained annotated genes (**Table 2**), a considerably higher proportion than identified by GPSM (between 30% and 34% across datasets). Of the 3,740 significant windows identified by RAiSD, 17 showed overlap with three different COJO associations in the young Red Angus dataset. Still, we did not observe any overlap with COJO SNPs in the full Red Angus dataset. In Red Angus, there were 12 sweep regions identified by both RAiSD and nSL. These included a signature on chromosome 6 (~78.95 Mb) that resides in a pleiotropic QTL for stayability, calving ease, and udder structure in dairy cattle, traits all under selection in the Red Angus breed (Cole et al. 2011). Another shared sweep region on chromosome 11 (18.1 Mb) is associated with multiple carcass quality traits (Mateescu et al. 2017).

**Table 2.** Number of significant windows within, proximal, or outside of genic regions identified by nSL and RAiSD.

| Breed | Analysis | Total Windows | Within Gene | Gene Proximal (< 50kb to nearest) | Intergenic |
| --- | --- | --- | --- | --- | --- |
| Red Angus | nSL | 365 | 183 (50%) | 73 (20%) | 109 (30%) |
| Red Angus | RAiSD | 6,502 | 3,453 (53%) | 811 (13%) | 2,238 (34%) |
| Simmental | nSL | 347 | 188 (54%) | 60 (17%) | 99 (29%) |
| Simmental | RAiSD | 6,497 | 3,030 (47%) | 1108 (17%) | 2,359 (36%) |

While the associated regions were vastly different from those identified by GPSM analysis, we identified twelve GPSM COJO SNPs (4 in full, 8 in young Red Angus datasets, respectively) that reside within 50 kb of a significant nSL window. Two of these loci reside on chromosome 14 (23.0 Mb and 58.1 Mb). The locus at 23.0 Mb was also identified as a significant sweep region by RAiSD. This locus resides within the gene *TMEM68,* which may be a driver of feed intake and growth phenotypes in cattle (Lindholm-Perry et al. 2012), and height in humans (Kichaev et al. 2019). This locus also resides within a QTL for Insulin-like growth factor 1 level (Fortes et al. 2013). The locus at 58.1 Mb lies within the gene Oxidation Resistance 1 (*OXR1*), which is also a known regulator of carcass weight in cattle (Zhang et al. 2019) and neutrophil counts in humans (Chen et al. 2020). A final shared region on chromosome 12 includes a region located immediately downstream of the gene *DNAJC15*, a heat shock protein with multiple reproductive associations in cattle (Zhang et al. 2011; Cochran et al. 2013) and human birth weight (Comuzzie et al. 2012).

We expect that selective sweeps assert different pressures on neighboring neutral sites compared with polygenic selection. To quantify these differences, we calculated linkage disequilibrium r^2^ statistics for all imputed SNPs within 100 kb of the lead SNP in significant RAiSD (n = 35) and nSL (n = 40) windows and significant GPSM COJO SNPs (n = 7) on chromosome 2 (**Figure 2E**). On average the r^2^ in regions surrounding sweep loci (RAiSD loci mean r^2^ = 0.223 [sd = 0.324], nSL loci mean r^2^ = 0.155 [sd = 0.205],) were significantly higher (Tukey HSD p-value < 2 × 10^−16^) than those around GPSM loci (mean r^2^ = 0.092 [sd = 0.192]).

Similarly, low levels of overlap between GPSM and sweep-detection methods existed in Simmental. A single GPSM COJO SNP resided within a significant RAiSD window (chromosome 26 at 637,478 base pairs). The lone overlap between nSL and GPSM COJO SNPs in Simmental was on chromosome 5 at 57,701,350 base pairs. This is approximately 500 kb from the *PMEL*/*ERBB3* coat color candidate locus and resides within a cluster of olfactory-associated genes. RAiSD and nSL analyses in Simmental identified at least 14 shared regions of selection (significant windows < 50 kb apart). This complementary evidence increases our confidence that an actual sweep occurred at a locus. The most notable shared locus between nSL and RAiSD is at the *POLLED* locus (Medugorac et al. 2012; Wiedemar et al. 2014) on chromosome 1, responsible for the presence or absence of horns. This locus has been under strong selection in most cattle populations and is frequently identified in selection mapping studies (Kemper et al. 2014; Xu et al. 2015). Another shared region of selection on chromosome 16 at ~42.6 Mb near the genes *MASP2* and *TARDBP* has been identified in numerous other sweep mapping studies of Simmental cattle (Ramey et al. 2013; Rothammer et al. 2013; Qanbari et al. 2014; Zhao et al. 2015).

While overlapping nSL and RAiSD signatures can help bolster confidence in candidate selective sweeps, most windows are uniquely identified by a single method (**Supplementary Tables 16-19**). For example, the locus with the most significant RAiSD μ statistic (**Supplementary Figures 5 & 6**) is between two major clusters of T cell receptors on chromosome 10 at ~24.2 Mb. This sweep region was also identified in European Simmental populations by Qanbari et al. (2014) and Zhao et al. (2015) (Qanbari et al. 2014; Zhao et al. 2015), illustrating the long-lasting signatures created by some selective sweeps. The next highest RAiSD μ statistic windows in Simmental are immediately upstream of *MC1R*, the gene responsible for red versus black coat color in cattle (Gutiérrez-Gil et al. 2007). The most significant nSL signature with a clear candidate gene resided within *TMEM132D*, a gene that has been identified in numerous selective sweep analyses (Qanbari et al. 2014; Mészáros et al. 2019; Moradian et al. 2020; Moscarelli et al. 2020) and is pleiotropic across dairy and beef cattle breeds (Martins et al. 2020).

## Discussion

This study further demonstrates the power of linear mixed models applied to a novel dependent variable for detecting ongoing selection in populations with temporal genotype data. The Generation Proxy Selection Mapping (GPSM) method (Decker et al. 2012; Rowan et al. 2021) is unique among selection mapping methods because it does not rely on outlier definitions, and significance is calculated on a per-marker basis, allowing us to pinpoint selection to very small intervals. Building on the work of Rowan et al. (2021), we expand GPSM to significantly larger datasets (46,454 and 78,787 animals) with imputed sequence level variants (> 11 million SNPs) (Rowan et al. 2021). This boost in power and resolution allowed us to map hundreds of small directional shifts in allele frequency, consistent with polygenic selection (Rosenberg et al. 2019). Further, by using a genomic relationship matrix, we can better control for the extensive population and family structure that exist in populations.

Due to the non-normality of generation proxy phenotypes in genotyped livestock populations, we explored the effects of variable transformation and data subsetting on GPSM’s ability to detect selection. Large residuals for individuals born in the distant past reduced the power to detect selection in the full Red Angus dataset. When subsetting this dataset to individuals born very recently or Box-Cox transforming the “birth date” generation proxy, we detected three times as many birth date-associated SNPs. In populations with low effective sizes (N_e_), we might expect stochastic allele frequency changes due to drift to generate detectable changes in frequency (Nei and Tajima 1981). Still, simulations performed in our previous work have shown that GPSM can effectively distinguish between drift and selection (Rowan et al. 2021). While a Box-Cox-transformed birth date generation proxy boosted the significance of signals in Red Angus, a similar optimal power transformation did not have the same effect when applied to Simmental animals. As a result, we used untransformed birth date as our dependent variable for sequence-level analyses but partitioned the data into subsets to probe different components of selection.

Identifying selection with SNPs on average 250 base pairs apart followed by conditional-joint analysis (COJO) allowed us to overlay epigenetic and open chromatin data generated by the Functional Annotation of Animal Genomes (FAANG) project onto the independently associated GPSM SNPs. We map hundreds of selected loci with predicted epigenetic marks in cattle (Giuffra et al. 2019). While GPSM COJO SNPs are not enriched in epigenetic mark regions genome-wide, the addition of these annotations allowed us to postulate the regulatory elements targeted by selection at particular GPSM loci. This selection on SNPs within epigenetic marks and other likely cis-regulatory regions suggests that selection on complex phenotypes alters gene expression in complex networks (Chan et al. 2011; Boyle et al. 2017; Liu et al. 2019).

While most other tests of polygenic selection explicitly test for correlations between allele frequencies and phenotypes over sampled time periods (Turchin et al. 2012; Beissinger et al. 2018; Szpiech et al. 2020), or between distinct diverged populations (Chen et al. 2010; Szpiech et al. 2020), GPSM operates agnostic to phenotype and population label. This expands the odds of detecting polygenic selection signatures but can make interpreting signals difficult. Fortunately, hundreds of QTL-mapping studies have been performed in cattle, providing an extensive database of loci known to affect economically important complex traits (Hu et al. 2019). We queried the Animal QTL Database for QTL near GPSM associations and performed enrichment tests to understand the traits under selection in these populations. In Red Angus, we identified significant QTL enrichments for several production traits such as body weight, average daily gain, and carcass weight, all traits that we would expect to find under selection in beef populations. We also identified multiple QTL classes influencing maternal traits and calving ease, two recent selection emphases in the Red Angus breed. By analyzing both the full Simmental dataset and a subset of purebred animals, we could disentangle which allele frequencies were changing due to Simmental-specific selection versus selection on variation introgressed from other breeds. In purebred Simmental, we found limited selection on traits that were not explicitly involved in appearance characteristics. Strong selection at the *PMEL*/*ERBB3* and *KIT* loci have been a significant focus of the breed as it aimed to make animals appear more like Angus cattle. As a result, GPSM detected less selection on complex production traits in the purebred dataset than the full dataset where we detected an excess of associations with carcass, production, and reproductive traits, consistent with ongoing selection in the cattle industry at large. Differences in the enriched carcass and production traits are consistent with traits that show appreciable average phenotypic differences between Angus and Simmental animals. While these QTL databases are far from comprehensive due to their biases towards loci with detectable effect sizes in frequently measured traits, they provide a valuable first step to identifying the production traits driving genomic changes in these populations. Further, we anticipate that GPSM loci will serve as useful functional annotations for the bovine genome. As studies continue to map associations with complex traits, knowledge of the ongoing selective forces acting upon those loci will add valuable context and information.

While some overlap between selective sweep mapping outlier windows (nSL and RAiSD) and GPSM hits existed, they largely identify different genomic loci. An intuitive explanation of this result is that selective sweeps remove variation from the locus; thus, there is little variation at that locus on which polygenic selection can act. Sweep mapping methods consistently identify important Mendelian loci, such as *POLLED*, coat color genes (*MC1R*, *KIT*, etc.), and large-effect QTL where selection has for all intents and purposes completed. GPSM’s strength is detecting subtle, directional shifts in allele frequency over short periods of ongoing selection. In contrast, nSL, RAiSD, and other sweep-mapping methods identify characteristics of sweeps such as extended haplotype homozygosity, long-range L.D., and changes to the local site frequency spectrum. We observed significantly less L.D. between GPSM SNPs and neighboring neutral sites compared with the lead SNPs in sweep peaks identified by nSL and RAiSD. Most sweep mapping strategies search for signatures at neighboring neutral sites, whereas GPSM tracks actual allele frequency changes over time.

In Red Angus and Simmental, we have identified the historical hard sweeps, soft sweeps, and ongoing polygenic selection that has altered the genotypes and phenotypes of these populations. Loci identified by RAiSD μ statistic have altered linkage disequilibrium, sequence diversity, and site frequency spectrum indicative of severe sweeps. In both Red Angus and Simmental, the genes identified by RAiSD were enriched for developmental processes, such as “anatomical structure development” and “embryonic skeletal system morphogenesis.” This likely reflects historical visual selection for “type” (body shape) and breed character (how closely an animal matches the breed’s ideal) that occurred before systematic data collection and mixed model genetic evaluations. Conversely, no functions were consistently enriched based on genes within nSL haplotype homozygosity regions, which signatures reflect weaker, incomplete sweeps. Genes identified by GPSM reflect genetic and genomic selection for economically important traits. This agrees with previous reports but with greater power and precision in the current study (Rowan et al. 2021). Genetic improvement in Red Angus is driven by selection within the breed, whereas improvement in Simmental appears to be caused by preferential introgression of certain Angus sires. Our results provide a detailed description of the selection architectures in two of the most common cattle breeds in America, reflecting a history of visual selection for breed type followed by a transition to modern genomic selection.

## Conclusions

Using large, commercially-generated cattle genotype datasets imputed to 11 million SNPs and the Generation Proxy Selection Mapping method, we fine-mapped hundreds of loci undergoing subtle directional shifts in frequency. These loci reside overwhelmingly in, or nearby, genes, suggesting that selection on complex traits is likely concerned with perturbing gene expression patterns in complex networks. GPSM detected largely different sets of selected loci than the selective sweep mapping methods. When longitudinally sampled genotypes are available, GPSM is a powerful method for detecting ongoing changes to the genome. This makes it a complementary approach to sweep-mapping strategies as we work to understand the impacts of all types of selection on the genome.

## Methods

### Genotype data and imputation

We used commercially generated assay genotypes from two populations of *Bos taurus* beef cattle. These data, made up of assays ranging in density from 25K to 777K SNPs, were filtered, phased, then imputed using the approach described in Rowan et al. (2019) (Rowan et al. 2019). Briefly, before phasing and imputation, we removed individuals with low call rates (< 0.90) and SNPs with low call rates (< 0.90) or extreme Hardy-Weinberg p-values (< 10^−50^, indicative of genotyping errors) in PLINK (version 1.9) (Purcell et al. 2007). Genotypes were phased using a high-density reference in Eagle v2.4.1 (Loh et al. 2016) and imputed using Minimac4 (Das et al. 2016). The resulting high-density chip-imputed dataset contained 811,967 autosomal SNPs for 90,580 Simmental and 46,454 Red Angus animals. We refer to this dataset as 811K throughout. High-density imputed genotypes were then imputed to 43,214,290 SNPs from whole-genome resequencing data using 9,871 reference individuals from the Thousand Bulls Project (Hayes and Daetwyler 2019). We restricted the imputation reference to high quality (Variant Quality Score Recalibration (McKenna et al. 2010) Tranche 90), biallelic variants with minor allele counts greater than 20. Following imputation, we further filtered imputed SNPs based on internally-calculated imputation R^2^ (> 0.4) and minor allele frequency (> 0.01), leaving 11,759,568 imputed sequence variants for downstream analysis. Genomic coordinates for all array and sequence genotypes were based on positions in the ARS-UCD1.2 assembly (Rosen et al. 2020).

### Generation Proxy Phenotypes

We use the breeder-reported birth date to calculate the continuous number of years (months and days expressed as a decimal) between the animal’s birth date and October 19, 2020. We used his value as the continuous generation proxy in GPSM. In addition, due to the extreme left-skewness of animal birth dates in the dataset, we performed a Box-Cox transformation of animal birth dates to test the effects of transforming the data for normality.

### Generation Proxy Selection Mapping (GPSM)

Generation proxy selection mapping uses an individual’s generation number or a proxy for generation number as the dependent variable in a genome-wide linear mixed model. To control for shared ancestry between individuals and to estimate variance components, we used autosomal SNP markers in our 811K imputed dataset with MAF > 0.01 to construct a genomic relationship matrix (GRM) with the method in Yang et al. (2011) (Yang et al. 2011) for each population. Variance components were estimated using a genomic restricted maximum-likelihood (GREML) approach implemented in GCTA (version 1.92.3) (Yang et al. 2010; Yang et al. 2011). To evaluate the impact of transformations to generation proxy phenotypes, we predicted random genetic effects (breeding values) and residuals for individuals using GCTA’s “--reml-pred-rand” function.

The model used for selection mapping was as follows:

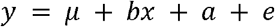

Here, □ is a vector of animal generation numbers or generation proxies, *μ* is the sample mean, *bx* is the scalar regression coefficient *b* on an N length vector of animal genotypes *x*, *a* is a random vector of polygenic terms 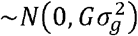 where *G* is a genomic relationship matrix, and *e* is a random error term 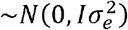. We performed all GPSM analyses using the “--mlma” function in GCTA. When testing the impact of generation proxy transformations on statistical power, we used 811K imputed genotypes. We performed GPSM on sequence-imputed genotypes on four total datasets with GRMs calculated with 811K genotypes.

To further refine GPSM signals to their core, we performed a within-analysis conditional and joint analysis (COJO) (Yang et al. 2012) in GCTA (v 1.92.3). COJO utilized summary data and genotypes from each of our sequence-level GPSM runs. We conditioned the COJO model on SNPs with GPSM p-values < 10^−5^. We controlled for SNP collinearity by setting conditional p-values of highly-correlated variants (r^2^ > 0.9) to 1. Significant COJO SNPs were those with conditional and genome-wide p-values < 5 × 10^−8^.

### Haplotype-based scans for selection

To map genomic regions that underwent strong selection in the distant-to-intermediate past, we used the number of segregating loci (nSL) method (Ferrer-Admetlla et al. 2014) on phased haplotypes from our full Red Angus and Simmental 811K datasets (MAF > 0.01), as well as on a subset of purebred Simmental animals. We implemented the nSL in selscan (Szpiech and Hernandez 2014) where per-SNP scores were normalized in 100 frequency bins. Instead of calculating significance in fixed windows, we fit a smoothing spline for each chromosome over normalized nSL scores using the GenWin R package (Beissinger et al. 2015). This allowed us to define variable-length windows in which we calculated the mean nSL scores. We considered the top 0.5% of these windows to be significant outliers for downstream annotation and analysis.

### Site frequency spectrum (SFS) scan for selection

We performed SFS-based scans for selection using called sequence genotypes from 14 American Red Angus, and 32 registered Simmental animals, some of which are crossbred, in the 1,000 Bull Genomes Project (Hayes and Daetwyler 2019) in RAiSD (v2.9) (Alachiotis and Pavlidis 2018). RAiSD calculates a composite selection statistic, μ, aimed at detecting various signatures of selective sweeps. To preserve the full site frequency spectrum, we did not filter on any quality or frequency-based metrics. Instead, we restricted our analysis to only biallelic SNPs. SNPs in the top 0.05% of RAiSD μ values were considered significant outliers (Alachiotis and Pavlidis 2018).

### Tajima’s D Scores

We used VCF-kit (Cook and Andersen 2017) to calculate Tajima’s D (Tajima 1989) scores across the genome for Red Angus animals in the 1,000 Bull Genomes project. We calculated these scores in fixed-width 100 kb bins across the genome.

### Gene & QTL annotation and enrichment

We annotated nearby genes and QTL using the GALLO R package with gene lists from ENSEMBL (Yates et al. 2020) and known *Bos taurus* QTL curated in the Animal QTL Database (Hu et al. 2019). We annotated all genes and QTL within 50 kb of significant (p < 5 × 10^−8^) COJO SNPs or sweep outlier regions (based on lead SNP). We performed QTL enrichment analysis with GALLO, using an FDR-corrected p-value threshold of 0.05 to identify traits whose known QTL were significantly enriched in GPSM signatures. We also annotated these SNPs with their Functional-And-Evolutionary Trait Heritability (FAETH) score as calculated in (Xiang et al. 2019), with coordinates lifted over from the UMD3.1.1 assembly to ARS-UCD1.2.

## Acknowledgments

We appreciate comments from Jeremy Taylor while writing this manuscript. We also appreciate farmers and ranchers for collecting this data and for data being shared by the Red Angus Association of America and the American Simmental Association. This work was supported by the Agriculture and Food Research Initiative (AFRI) grant No. 2016-68004-24827 from the U.S. Department of Agriculture (USDA) National Institute of Food and Agriculture (NIFA).

---

**Supplementary Table 1.** Genomic restricted maximum likelihood (GREML) estimates of proportion variance explained (PVE) for various statistical transformations to birth date used as generation proxy in Red Angus.

| Subset | n(animals) | Dependent Variable | PVE (S.E.) |
| --- | --- | --- | --- |
| FULL | 46,454 | Birth date | 0.523 (0.007) |
| FULL | 46,456 | Birth date<sup>-0.237</sup> | 0.657 (0.006) |
| Young Only<sup>1</sup> | 44,470 | Birth date | 0.773 (0.004) |
| Old Only<sup>2</sup> | 1,984 | Birth date | 0.557 (0.031) |

<sup>1</sup> Animals born on or after January 1, 2012
<sup>2</sup> Animals born prior to January 1, 2012

**Supplementary Table 2.**
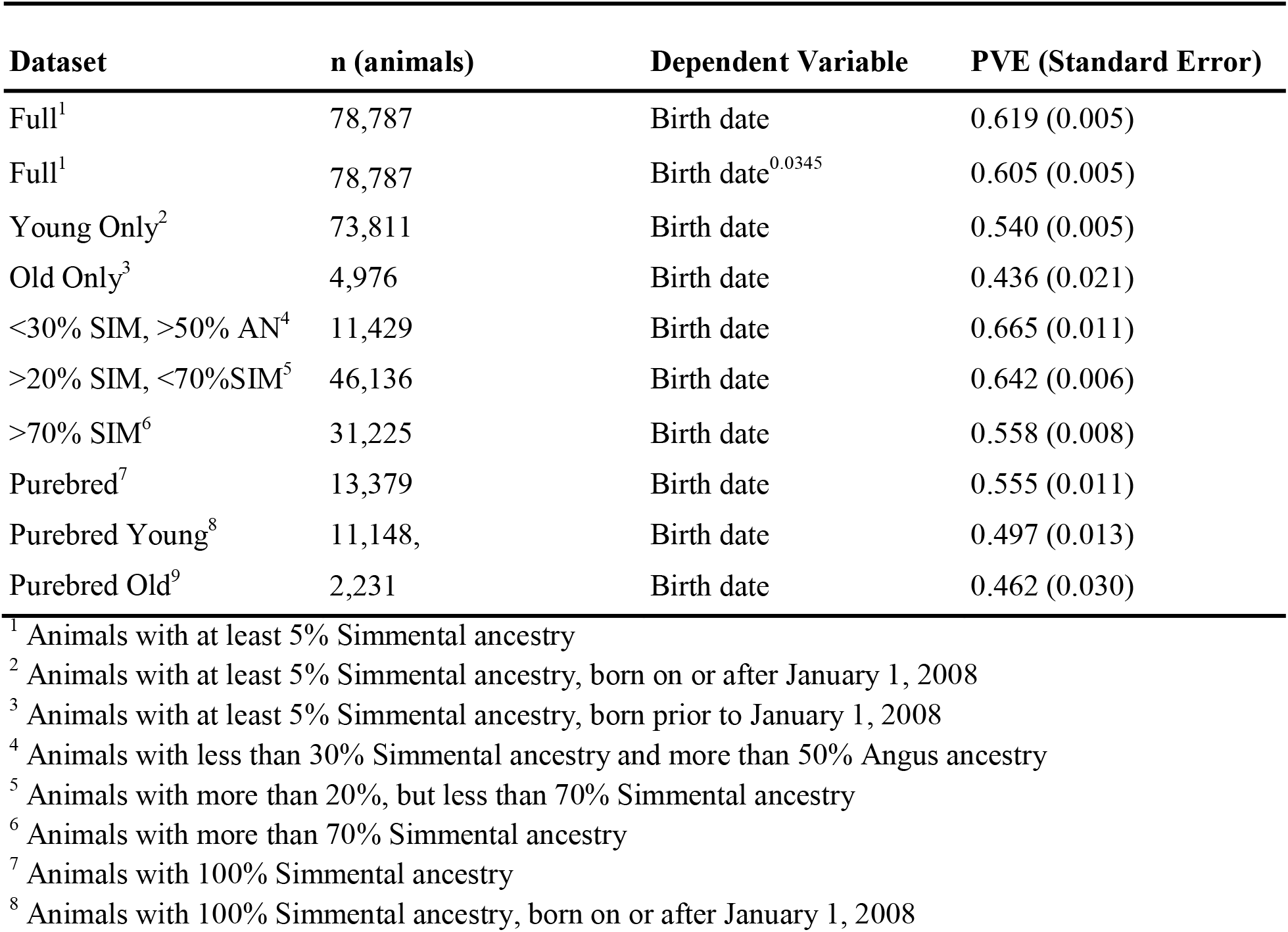
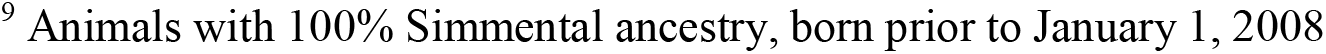
Genomic restricted maximum likelihood (GREML) estimates of proportion variance explained (PVE) for various subsets of the Simmental dataset.

| Dataset | n (animals) | Dependent Variable | PVE (Standard Error) |
| --- | --- | --- | --- |
| Full<sup>1</sup> | 78,787 | Birth date | 0.619 (0.005) |
| Full<sup>1</sup> | 78,787 | Birth date<sup>0.0345</sup> | 0.605 (0.005) |
| Young Only<sup>2</sup> | 73,811 | Birth date | 0.540 (0.005) |
| Old Only<sup>3</sup> | 4,976 | Birth date | 0.436 (0.021) |
| <30% SIM, >50% AN<sup>4</sup> | 11,429 | Birth date | 0.665 (0.011) |
| >20% SIM, <70%SIM<sup>5</sup> | 46,136 | Birth date | 0.642 (0.006) |
| >70% SIM<sup>6</sup> | 31,225 | Birth date | 0.558 (0.008) |
| Purebred<sup>7</sup> | 13,379 | Birth date | 0.555 (0.011) |
| Purebred Young<sup>8</sup> | 11,148, | Birth date | 0.497 (0.013) |
| Purebred Old<sup>9</sup> | 2,231 | Birth date | 0.462 (0.030) |

<sup>1</sup> Animals with at least 5% Simmental ancestry
<sup>2</sup> Animals with at least 5% Simmental ancestry, born on or after January 1, 2008
<sup>3</sup> Animals with at least 5% Simmental ancestry, born prior to January 1, 2008
<sup>4</sup> Animals with less than 30% Simmental ancestry and more than 50% Angus ancestry
<sup>5</sup> Animals with more than 20%, but less than 70% Simmental ancestry
<sup>6</sup> Animals with more than 70% Simmental ancestry
<sup>7</sup> Animals with 100% Simmental ancestry
<sup>8</sup> Animals with 100% Simmental ancestry, born on or after January 1, 2008
<sup>9</sup> Animals with 100% Simmental ancestry, born prior to January 1, 2008

**Supplementary Table 3.**
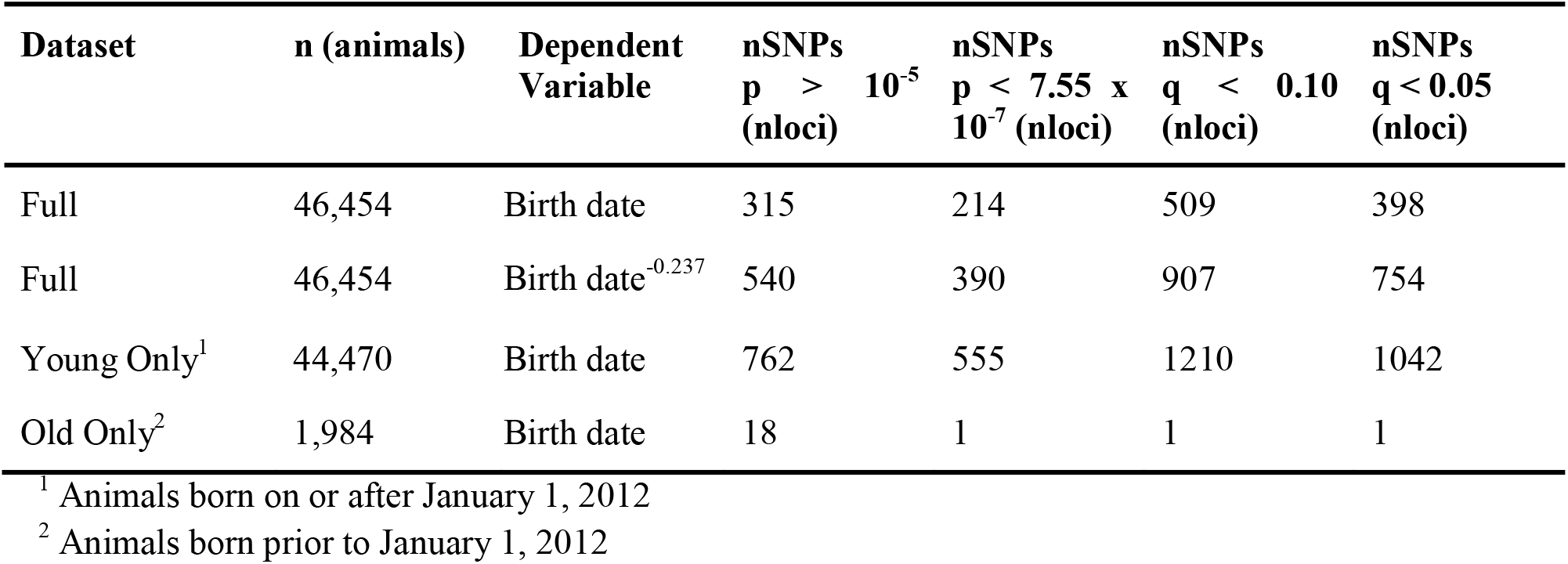
Counts of significant SNPs identified in each GPSM analysis of the Red Angus population using different transformations to generation proxy as dependent variable in 811K SNPs. The four significance cutoffs are reported: 1) A nominal significance (p < 10^−5^), 2) a Bonferroni-adjusted threshold (p < 7.55 × 10^−7^), and FDR-corrected q-values 3) < 0.10 or 4) < 0.05.

**Supplementary Table 4.** Counts of significant SNPs identified in each GPSM analysis of the Simmental population using different population subsets and transformations to generation proxy with 811K SNPs. Ancestry proportions are pedigree estimates reported by American Simmental Association. The four significance thresholds are reported: 1) A nominal significance (p < 10^−5^), 2) a Bonferroni-adjusted threshold (p < 7.55 × 10^−7^), and FDR-corrected q-values 3) < 0.10 or 4) < 0.05.

| Dataset | n (animals) | Dependent Variable | Nominal<br>($p < 10^{-5}$) | Bonferroni | $q < 0.1$ | $q < 0.05$ |
| --- | --- | --- | --- | --- | --- | --- |
| Full<sup>1</sup> | 78,787 | Birth date | 120 | 70 | 137 | 117 |
| Full<sup>1</sup> | 78,787 | Birth date<sup>0.0345</sup> | 109 | 77 | 130 | 105 |
| Young Only<sup>2</sup> | 73,811 | Birth date | 94 | 68 | 100 | 89 |
| Old Only<sup>3</sup> | 4,976 | Birth date | 119 | 72 | 171 | 115 |
| <30% SIM, >50% AN<sup>4</sup> | 11,429 | Birth date | 14 | 3 | 3 | 0 |
| >20% SIM, <70% SIM<sup>5</sup> | 46,136 | Birth date | 46 | 31 | 39 | 38 |
| >70% SIM<sup>6</sup> | 31,225 | Birth date | 100 | 61 | 107 | 88 |
| >70% SIM<sup>6</sup> | 31,225 | □□□ (□□□□□ □□□ | 51 | 24 | 44 | 32 |
| Purebred<sup>7</sup> | 13,379 | Birth date | 92 | 50 | 111 | 85 |
| Purebred Young<sup>8</sup> | 11,148, | Birth date | 11 | 4 | 4 | 3 |
| Purebred Old<sup>9</sup> | 2,231 | Birth date | 60 | 34 | 59 | 48 |

<sup>1</sup> Animals with at least 5% Simmental ancestry
<sup>2</sup> Animals with at least 5% Simmental ancestry, born on or after January 1, 2008
<sup>3</sup> Animals with at least 5% Simmental ancestry, born prior to January 1, 2008
<sup>4</sup> Animals with less than 30% Simmental ancestry and more than 50% Angus ancestry
<sup>5</sup> Animals with more than 20%, but less than 70% Simmental ancestry
<sup>6</sup> Animals with more than 70% Simmental ancestry
<sup>7</sup> Animals with 100% Simmental ancestry
<sup>8</sup> Animals with 100% Simmental ancestry, born on or after January 1, 2008
<sup>9</sup> Animals with 100% Simmental ancestry, born prior to January 1, 2008

**Supplementary Tables 5-16 are located in the associated Excel file.**

**Supplementary Table 5.** Significant (p < 5 × 10^−8^) conditional and joint (COJO) SNPs from full Red Angus GPSM analysis and their annotated nearby genes (< 50 kb).

**Supplementary Table 6.** Significant (p < 5 × 10^−8^) conditional and joint (COJO) SNPs from young Red Angus GPSM analysis and their annotated nearby genes (< 50 kb).

**Supplementary Table 7.** Significantly enriched QTL in regions (< 50 kb) from significant GPSM COJO SNPs identified in the full Red Angus dataset (n SNPs = 248).

**Supplementary Table 8.** Significantly enriched QTL in regions (< 50 kb) from significant GPSM COJO SNPs identified in the young Red Angus dataset (n SNPs = 417).

**Supplementary Table 9**. Significantly enriched gene set annotations in regions (< 50 kb) from significant GPSM COJO SNPs identified in the young Red Angus dataset (n SNPs = 417).

**Supplementary Table 10.** Significant (p < 5 × 10^−8^) conditional and joint (COJO) SNPs from full Simmental GPSM analysis and their annotated nearby genes (< 50 kb).

**Supplementary Table 11.** Significant (p < 5 × 10^−8^) conditional and joint (COJO) SNPs from purebred Simmental GPSM analysis and their annotated nearby genes (< 50 kb).

**Supplementary Table 12.** Significantly enriched QTL in regions (< 50 kb) from significant GPSM COJO SNPs identified in full Simmental dataset (n SNPs = 344).

**Supplementary Table 13.** Significantly enriched QTL in regions (< 50 kb) from significant GPSM COJO SNPs identified in purebred Simmental dataset (n SNPs = 33).

**Supplementary Table 14**. Significantly enriched gene set annotations in regions (< 50 kb) from significant GPSM COJO SNPs identified in the full Simmental dataset (n SNPs = 344).

**Supplementary Table 15.** Significantly enriched gene set annotations in regions (< 50 kb) from significant GPSM COJO SNPs identified in the purebred Simmental dataset (n SNPs = 33).

**Supplementary Table 16.** Red Angus outlier nSL windows (top 0.5%) as defined by GenWin R package. Genes that fell within significant windows are reported with their corresponding window.

**Supplementary Table 17.** Unique genes within Red Angus outlier RAiSD windows (top 0.05%). Reported window is the window with the highest μ value that overlaps with the gene.

**Supplementary Table 18.** Simmental outlier nSL windows (top 0.5%) as defined by GenWin R package. Genes that fell within significant windows are reported with their corresponding window.

**Supplementary Table 19.** Unique genes within Simmental outlier RAiSD windows (top 0.05%). Reported window is the window with the highest μ value that overlaps with the gene.

**Supplementary Table 20.** Significantly enriched gene set annotations of genes within Simmental outlier nSL windows.

**Supplementary Table 21.** Significantly enriched gene set annotations of genes within Red Angus outlier RAiSD windows.

**Supplementary Table 22.** Significantly enriched gene set annotations of genes within Simmental outlier RAiSD windows.

**Supplementary Figure 1.**
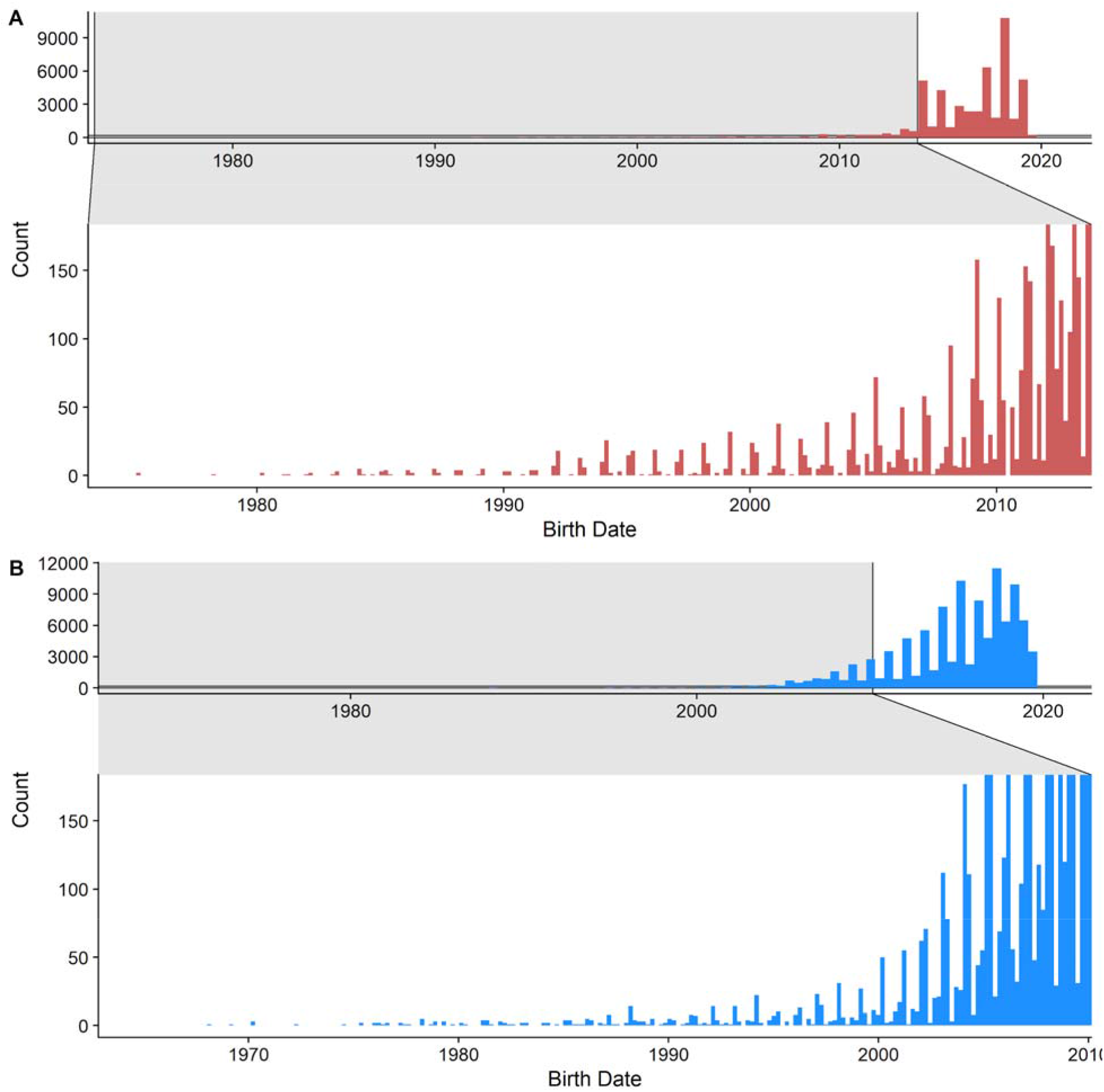
Birth date distributions of Red Angus and Simmental datasets. Histograms of (A) Red Angus and (B) Simmental datasets. Left tails of the distributions (older animals) have been zoomed in the lower half of each panel.

**Supplementary Figure 2.**
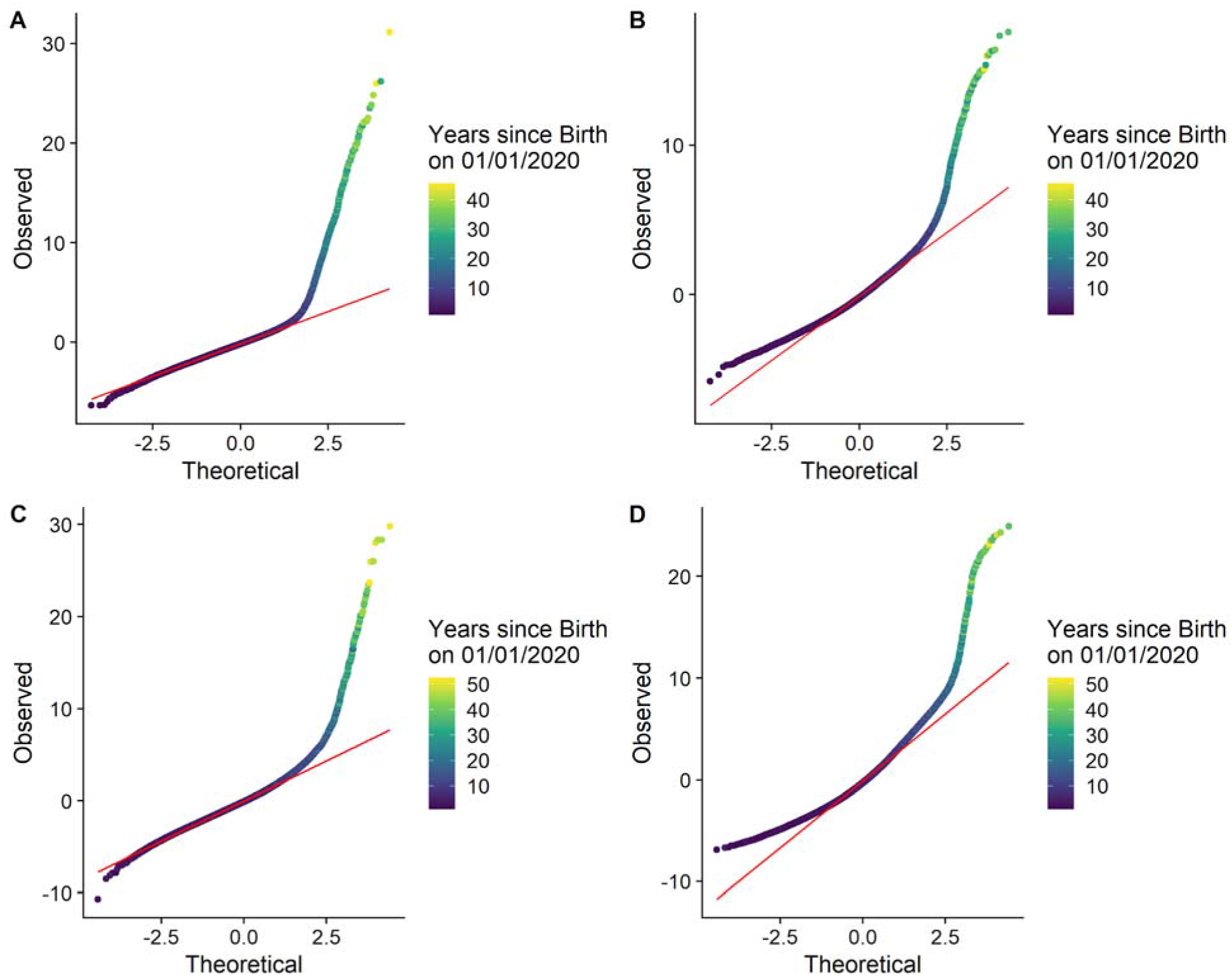
Q-Q plots from Birth Date GREML analysis. Q-Q plots of GREML (A) residuals and (B) estimated breeding values for Red Angus individuals. Each point represents an animal, colored by the animal’s age as of January 1st, 2020. Simmental GREML Q-Q plots of (C) residuals and (D) estimated breeding values.

**Supplementary Figure 3.**
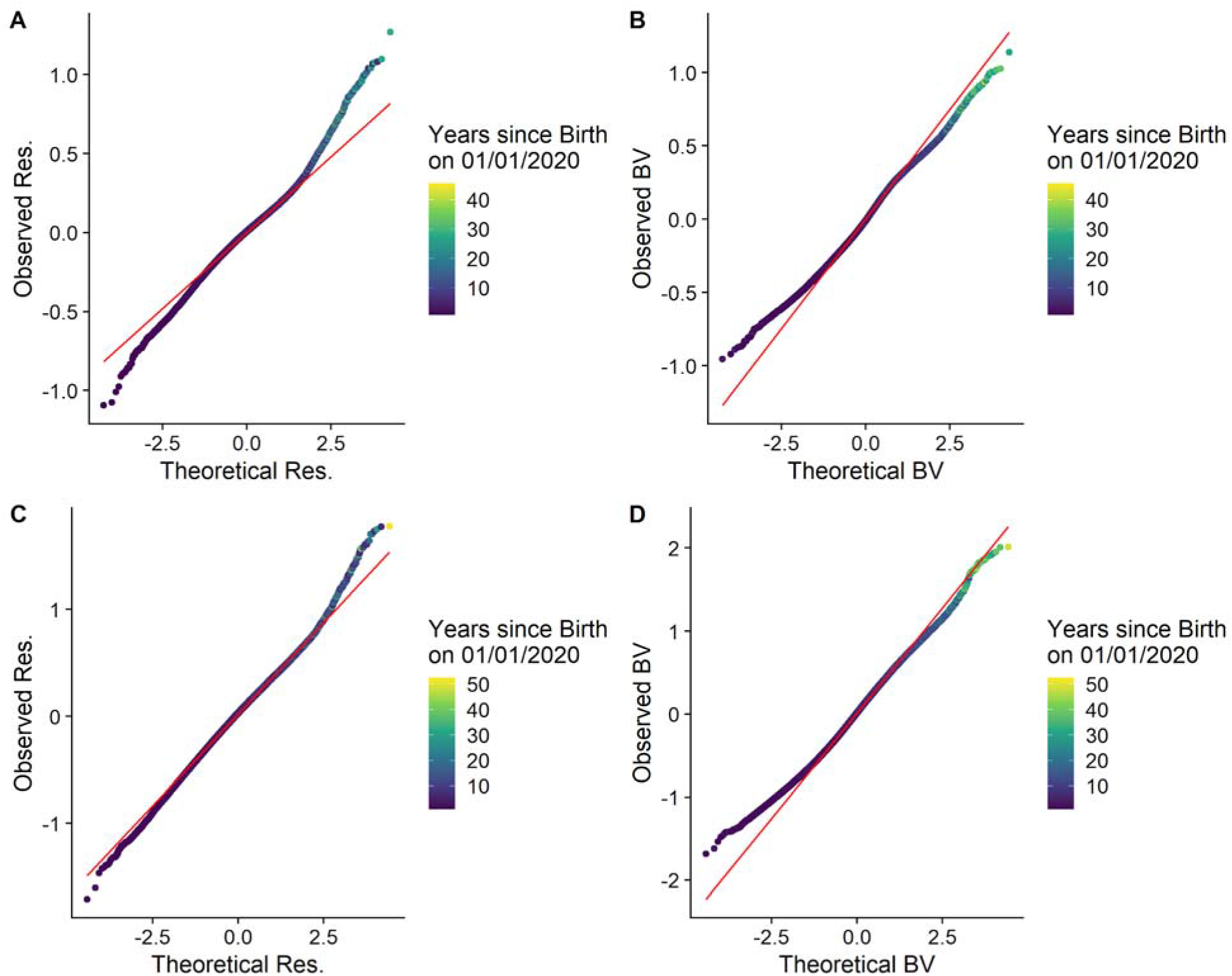
Q-Q plots from Box-Cox Transformed Birth Date GREML analysis. Q-Q plots of GREML (A) residuals and (B) estimated breeding values for Red Angus individuals using Box-Cox transformed birth date as the dependent variable. Each point represents an animal, colored by the animal’s age as of January 1st, 2020. Simmental GREML Q-Q plots of (C) residuals and (D) estimated breeding values from analysis using Box-Cox transformed birth date as dependent variable.

**Supplementary Figure 4.**
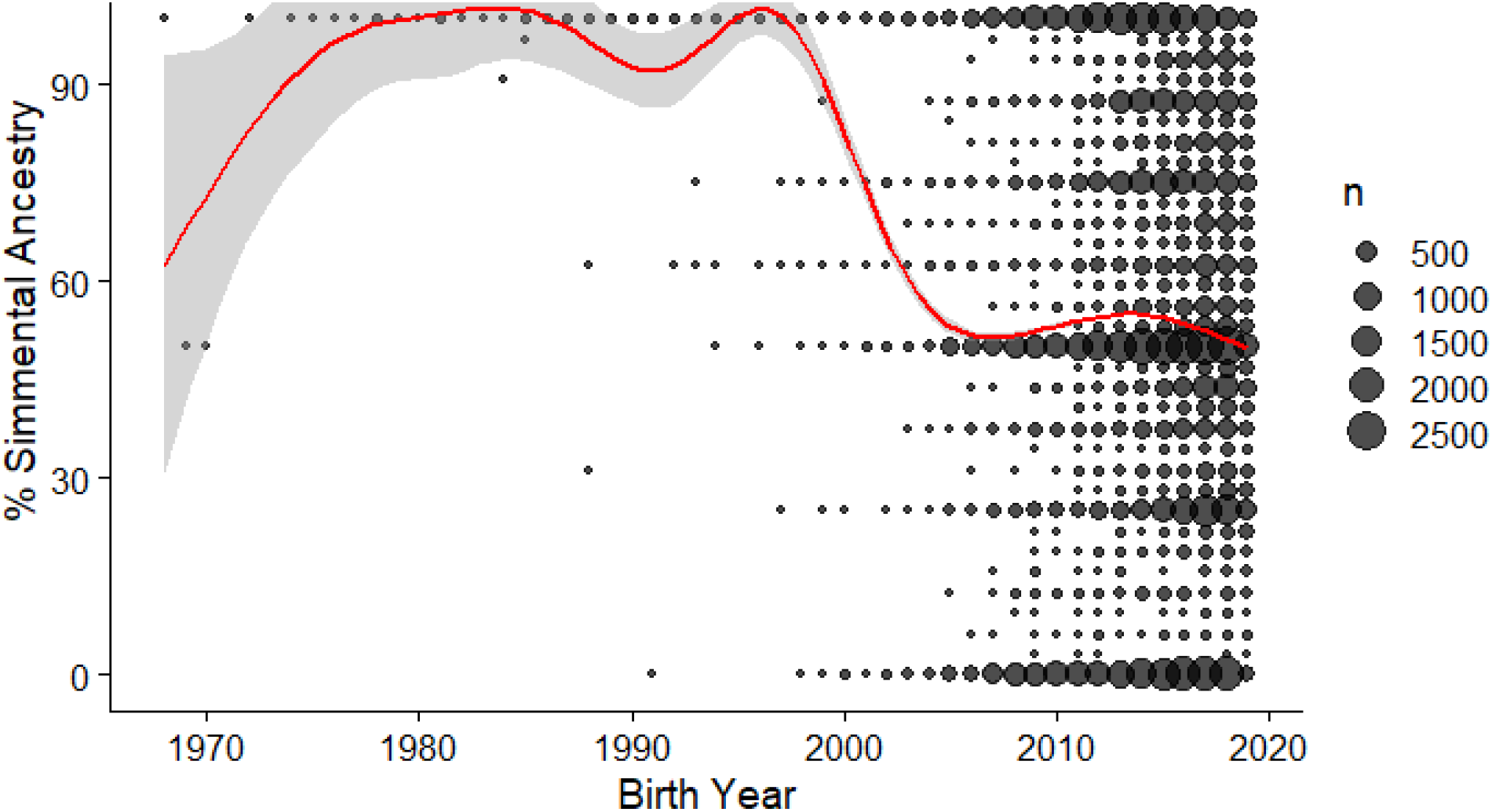
The changing breed composition of registered Simmental. Counts of Simmental ancestry percentages over time in all genotyped animals in the American Simmental dataset. Points represent birth year/% Simmental ancestry combinations in the data, sized by the number of animals in each of those classes. The red line is a smoothed mean, surrounded by a 95% confidence interval in grey.

**Supplementary Figure 5.**
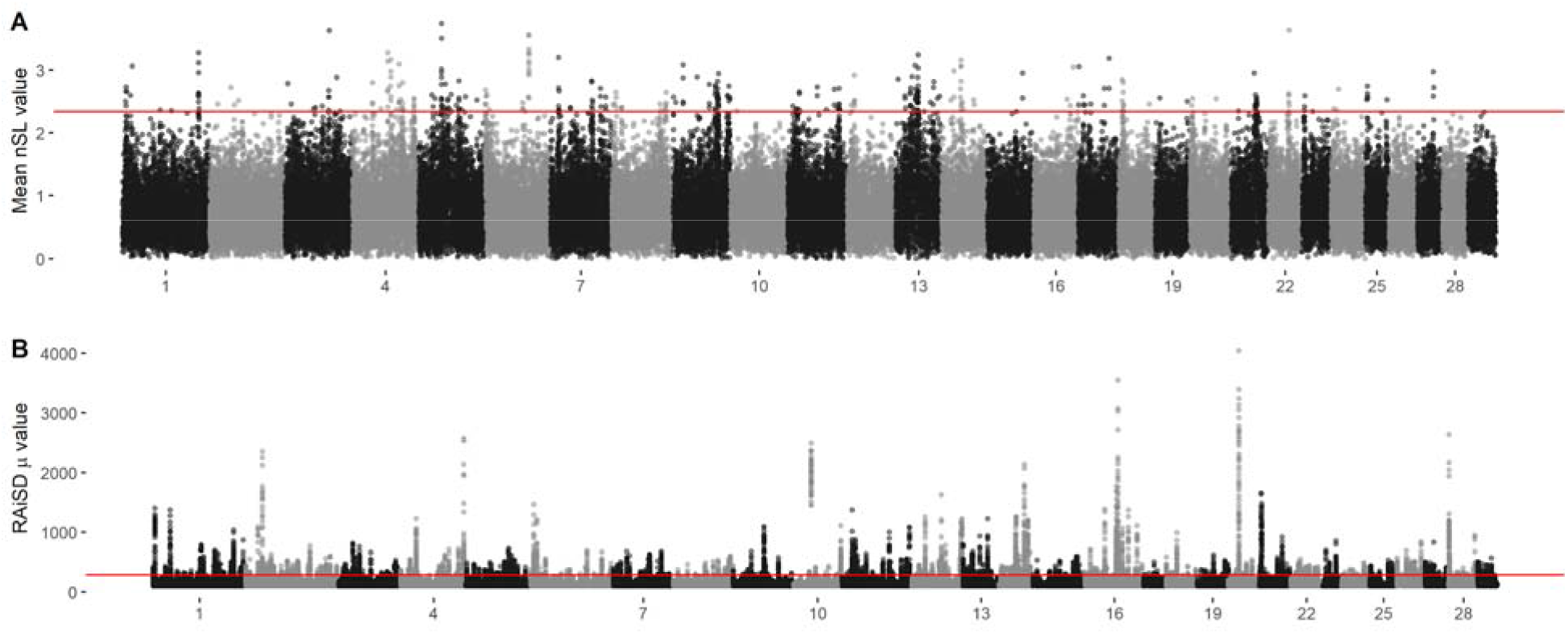
Selective sweep mapping in Red Angus. (A) Number of segregating loci (nSL) statistic windows across the genome. Window boundaries were defined by GenWin R package. Points represent the average |nSL| values within a window, with the genomic position defined as the genomic center of the window. Red line delineates 0.5% outliers deemed significant. (B) Manhattan plot of RAiSD μ statistics calculated from sequenced American Red Angus animals in the 1000 Bull Genomes Project. Red line delineates 0.05% outliers.

**Supplementary Figure 6.**
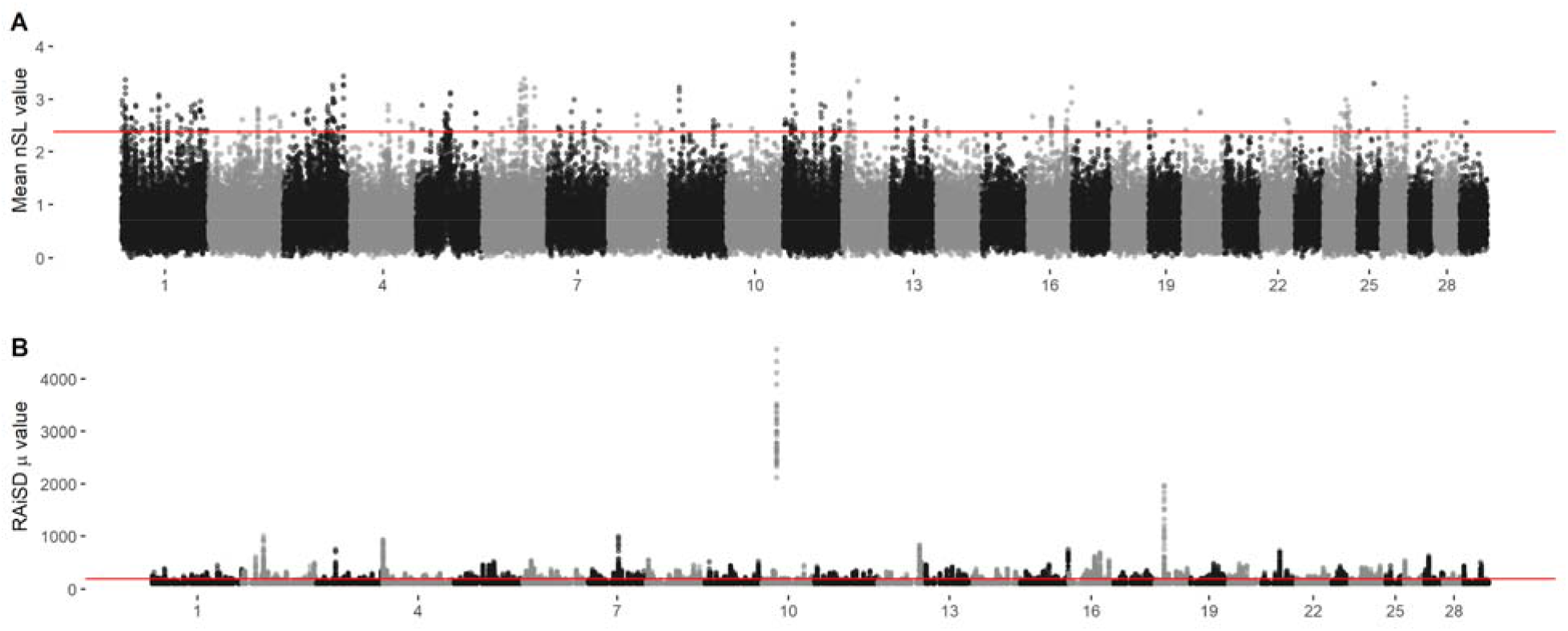
Selective sweep mapping in Simmental. (A) Number of segregating loci (nSL) statistic windows across the genome. Window boundaries were defined by the GenWin R package. Points represent the average |nSL| values within a window, with the genomic position defined as the genomic center of the window. Red line delineates 0.5% outliers deemed significant. (B) Manhattan plot of RAiSD μ statistic calculated from sequenced American Simmental animals in the 1000 Bull Genomes Project. Red line delineates 0.05% outliers.

